# Quantitative proteomic mass spectrometry of protein kinases to determine dynamic heterogeneity of the human kinome

**DOI:** 10.1101/2024.10.04.614143

**Authors:** Michael P. East, Robert W. Sprung, Denis O. Okumu, J. Felix Olivares-Quintero, Chinmaya U. Joisa, Xin Chen, Qiang Zhang, Petra Erdmann-Gilmore, Yiling Mi, Noah Sciaky, James P. Malone, Sonam Bhatia, Ian C. McCabe, Yi Xu, Matthew D. Sutcliffe, Jingqin Luo, Patricia A. Spears, Charles M. Perou, H. Shelton Earp, Lisa A. Carey, Jen Jen Yeh, David L. Spector, Shawn M. Gomez, Philip M. Spanheimer, R. Reid Townsend, Gary L. Johnson

## Abstract

The kinome is a dynamic system of kinases regulating signaling networks in cells and dysfunction of protein kinases contributes to many diseases. Regulation of the protein expression of kinases alters cellular responses to environmental changes and perturbations. We configured a library of 672 proteotypic peptides to quantify >300 kinases in a single LC-MS experiment using ten micrograms protein from human tissues including biopsies. This enables absolute quantitation of kinase protein abundance at attomole-femtomole expression levels, requiring no kinase enrichment and less than ten micrograms of starting protein from flash-frozen and formalin fixed paraffin embedded tissues. Breast cancer biopsies, organoids, and cell lines were analyzed using the SureQuant method, demonstrating the heterogeneity of kinase protein expression across and within breast cancer clinical subtypes. Kinome quantitation was coupled with nanoscale phosphoproteomics, providing a feasible method for novel clinical diagnosis and understanding of patient kinome responses to treatment.

## Introduction

The human kinome consists of ∼535 protein and lipid kinases integrated into cellular signaling networks for the control of many cellular functions and is frequently dysregulated in different diseases^1-4^. In many cancers, changes in expression or mutation of different protein kinases drive the cancer phenotype^5^. Because protein kinases are critically important for control of diverse cellular functions, their enzymatic phosphorylation of substrates is tightly controlled. The regulation of a kinase’s activity may involve protein-protein interactions, subcellular localization, allosteric small molecules (e.g., cAMP) and phosphorylation-dephosphorylation in addition to other covalent modifications^6-8^. Most human cells express 350-400 protein kinases and there is significant heterogeneity in the protein expression level of specific kinases in cells even of the same genotype and phenotype^9^. Furthermore, the kinome is dynamic with changes in the proteomic expression of kinases in response to cellular perturbations^10-14^.

The plasticity of the kinome is readily observed in the adaptive reprogramming of kinase expression in response to targeted kinase inhibitors^12-17^. For example, inhibition of mitogen activated protein kinase kinase 1 and 2 (MAP2K1/2 or MEK1/2) inhibits MEK1/2 phosphorylation and activation of extracellular-signal regulated kinase 1 and 2 (ERK1/2 or MAPK3/1)^13^. ERK1/2 inhibition results in c-Myc degradation and chromatin remodeling, causing transcriptional changes that include both tyrosine and serine/threonine protein kinases^11^. Inhibitor induced adaptive reprogramming is unique in the transcriptional and proteomic profile for each targeted kinase, and is seen with tyrosine kinase and serine/threonine kinase inhibitors^10-12^ and RAS inhibitors^18^, where RAS proteins are upstream activators of the MEK1/2-ERK1/2 signaling network^19^. A functional consequence of adaptive reprogramming is the proteomic landscape of the kinome is altered, resulting in new kinase signaling networks that frequently overcome the targeted kinase inhibition leading to resistance of tumor cells to therapy^17^. Given that there are currently 87 small molecule inhibitors^20^ and 10 antibodies that are FDA approved targeting specific protein kinases, quantitative proteomic measurements of the kinome and how kinase expression changes with environmental and pharmacological perturbations is required to understand the plasticity of kinome adaptation and altered cellular signaling.

Canonical bottom-up shotgun proteomics workflows fail to provide comprehensive coverage of the human kinome largely due to generally low levels of kinase expression in most cells. Affinity capture of kinases from cell lysates using broadly specific ATP-competitive kinase inhibitors immobilized on beads, called Multiplexed Inhibitor Beads (MIBs) and Kinobeads, drastically improved quantitation by mass spectrometry (MS)^13,21^. Analysis of functional kinase protein levels using MIB/MS revealed heterogeneity in expression across intrinsic breast cancer subtypes and was sufficient to distinguish between clinical subtypes in diagnostic patient tumor biopsies^10^. Probing the functional kinome with MIB/MS also established the dynamic nature of the kinome in response to targeted kinase inhibitors^10,13,15^. Although MIB/MS and Kinobeads have proven invaluable in our understanding of the behavior of the kinome, their utility in the clinical setting is limited by the material requirement of nondenatured proteins (∼1-2 mg of lysate protein), affinity of kinases for immobilized inhibitors, and capacity for only relative levels of quantitation from shotgun proteomics.

Parallel reaction monitoring (PRM) proteomics offers more quantitative and sensitive analysis of protein abundance^22^. This targeted method uses stable isotopically labeled (SIL) peptides spiked into tryptic digests from samples at known concentrations. Endogenous peptide abundance is determined by the relative intensity of MS2 product ions to matching product ions from SIL peptides. The SureQuant triggered acquisition software has further empowered this methodology by minimizing unproductive scan times and removing the need for retention time scheduling ultimately leading to better overall sensitivity^23^. We have designed and validated a SIL peptide library targeting ∼2/3 of the human protein kinome. SIL peptide performance was analyzed based on the guidelines set forth by the NCI’s Clinical Proteomic Tumor Analysis Consortium (CPTAC) for reproducibility, linearity of quantitation, and sensitivity. SureQuant/PRM MS analysis allows quantitative measure of dynamic and heterogenous kinase expression in patient tumor biopsies, formalin fixed paraffin embedded tissues (FFPE), organoids, and cell lines using 100-fold less starting material (10μg of protein or 50,000 - 100,000 cells) without kinase enrichment. The human kinase PRM library allows quantitative proteomic kinome measurements of clinical samples and is easily integrated with phosphoproteomics.

## Results

### Construction and validation of the kinome targeted SIL peptide library

Selection of proteotypic peptides for targeted proteomics analysis is challenging and remains largely empirical. Initial selection of candidate peptides based on standard criteria^24^ including tryptic peptide length (7-20 amino acids), efficiency of tryptic peptide production and lack of residues subject to variable modification resulting in distribution of signal over multiple m/z values (e.g. methionine, tryptophan, glutamine, asparagine, histidine) facilitates reproducible measurement of target peptides and the development of robust standards for quantitation. Despite these considerations, empirical approaches are necessary for optimal peptide assay development to maximize selectivity, sensitivity, precision, and minimize precursor concurrency. To guide selection of peptides, we used a complement of kinase inhibitors immobilized on Sepharose beads (MIBs) to capture kinases from patient tumor biopsies, xenografts and cell lines followed by data-dependent mass spectrometry (Fig. 1A)^13^. MIB/MS allows for the capture and MS analysis of >90% of the human kinome, providing the basis for selection of optimal proteotypic peptides for human kinases. This library of proteotypic candidates was further refined based on observed retention time during reverse phase liquid chromatography to minimize concurrency during LC-MS/MS analysis.

**Figure 1:**
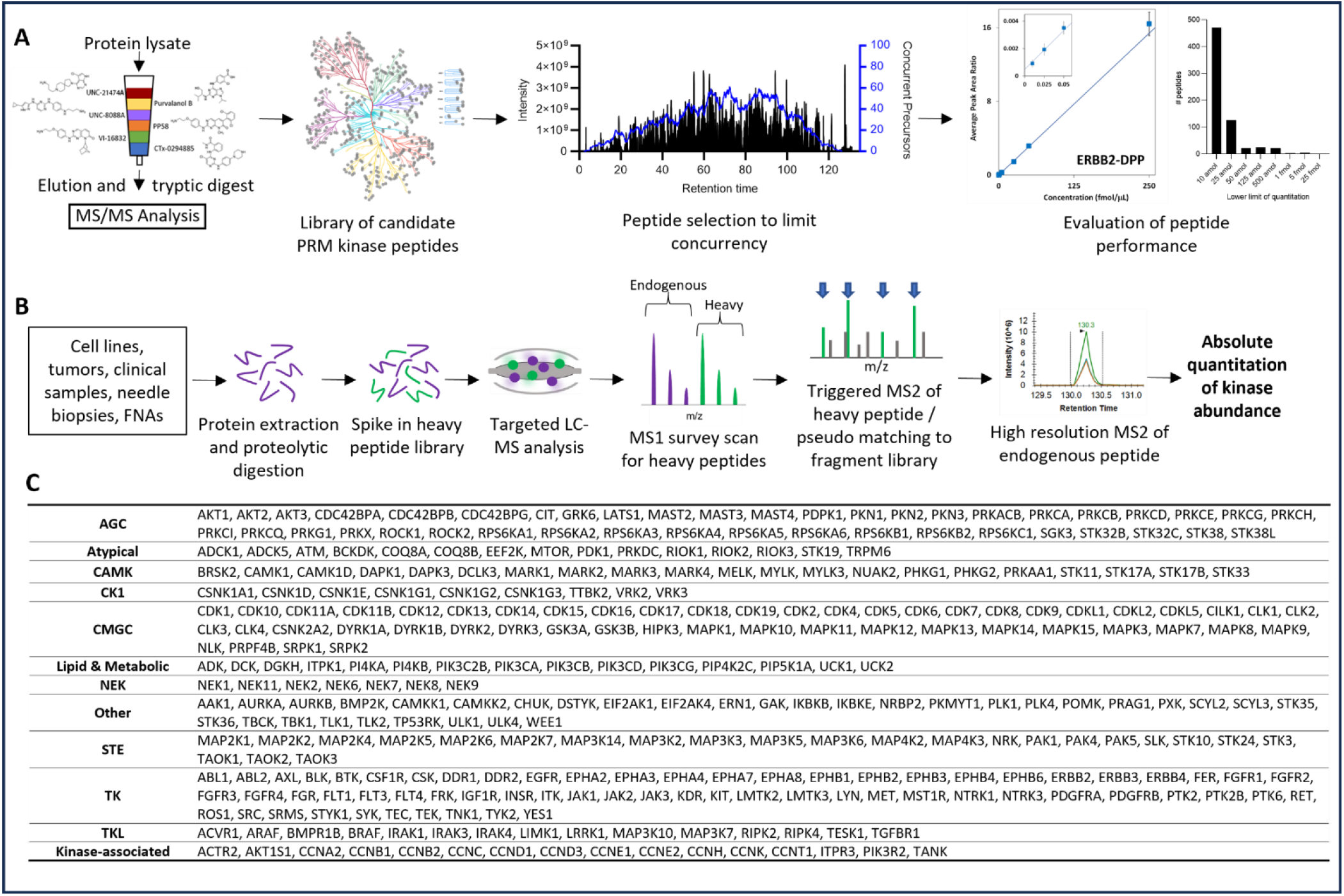
SureQuant triggered-parallel reaction monitoring library construction and workflow. **A)** Candidate proteotypic peptides for stable isotopically labeled (SIL) internal standards were identified empirically by kinase enrichment via multiplexed inhibitor beads (MIBs) coupled with MS/MS analysis^13^. The list of SIL candidates was refined based on peptide retention time to limit concurrency. SIL performance was validated based on Clinical Proteomic Tumor Analysis Consortium guidelines with lower limit of quantitation determined by calibration curves consisting of nine concentrations ranging from 10 amol to 250 fmol. **B)** SILs were spiked into sample tryptic digests at a concentration of 25 fmol/ug total protein prior to solid phase extraction of peptides. Internal standard triggered parallel reaction monitoring was performed using the Thermo SureQuant triggering software. Optimal charge states and product ions were determined by a survey run of SILs alone. SureQuant triggering software continuously scans for m/z ratios corresponding to SILs during MS1 then by pseudomatching product ions during MS2 to a defined list determined during the survey run. Upon pseudomatching at least three product ions, high-resolution MS2 scans for endogenous peptide are triggered and protein abundance quantified based on the intensity of the three most abundant product ions using the Skyline software package^70^. **C)** SureQuant/PRM library characterized to date consists of 672 peptides targeting 296 kinases, 16 kinase-associated proteins, and the “proteomic ruler” PARK7^37^.

Stable isotope labeled (SIL) peptides were synthesized with uniformly ^13^C and ^15^N labeled C-terminal amino acids. Performance in response curves and in intra- and inter-day reproducibility experiments were evaluated based on experimental guidelines established by the CPTAC^25^. The lower limit of quantitation and linearity of response for each SIL peptide was determined using forward response curves with heavy peptide abundance held constant at 25 fmol/injection over ten light isotope peptide amounts ranging from 10 amol to 250 fmol in triplicate on the Exploris 480 mass spectrometer using the SureQuant triggered acquisition parameters used for biological samples (Fig. 1B, Suppl Table 1, Suppl Fig. 1). For these assays, lower limit of quantitation was determined based on the concentration of analyte that could be measured with a coefficient of variance <30% using triplicate analyses. Using this criterion, 664 peptides had a LLOQ less than 1 fmol, representing 98.8% of the peptides assayed, enabling sensitive and reproducible quantitative standards for 313 kinases and kinase associated proteins (Fig. 1C, Extended Data Figure 1, Suppl Table 1)^26,27^.

### Quantitative measure of protein kinases in patient tumors

Kinome SureQuant/PRM offers unparalleled, absolute quantitation at attomole to femtomole kinase protein expression levels with no enrichment needed from clinical samples. Using a panel of six breast cancer patient tumors, approximately 190 kinases were quantified in one LC-MS analysis for each tumor (Suppl Table 2). The analysis gave quantitative measures of kinase families including CDKs (Fig. 2A, B) and RTKs (Fig. 2C, D). The SIL library targeted 18 of the 20 human CDKs (Fig. 2A). Peptides initially characterized for CDK20 and CDK3 were inadequate for quantitation and new SIL assays for these kinases are currently in development. Of the 18 targeted CDKs, 16 (including CDK11A/B) were quantified across breast cancer patient tumors, while endogenous peptides for CDK1 and 7 were not quantifiable due to low abundance or interference from unrelated product ions. Cell cycle CDKs include CDK1-6 and CDK14-18 whereas the transcriptional CDKs are comprised of CDK7-13 and CDK19-20^28^. While expression of CDK15, CDK16 and CDK17 was similar across patient tumors, protein expression for other CDKs was heterogeneous within the same SureQuant/PRM LC-MS analysis. For example, CDK9, the catalytic subunit of the P-TEFb complex responsible for allowing RNA-Pol II transcription elongation has been a target for development of small molecule therapeutic inhibitors^29^. Other transcriptional regulatory CDKs that show variation in expression with patient biopsies include CDK11A/B and CDK19, suggesting quantitative protein expression levels of transcriptional regulatory CDKs may be an important determinant for targeted inhibitor efficacy.

**Figure 2:**
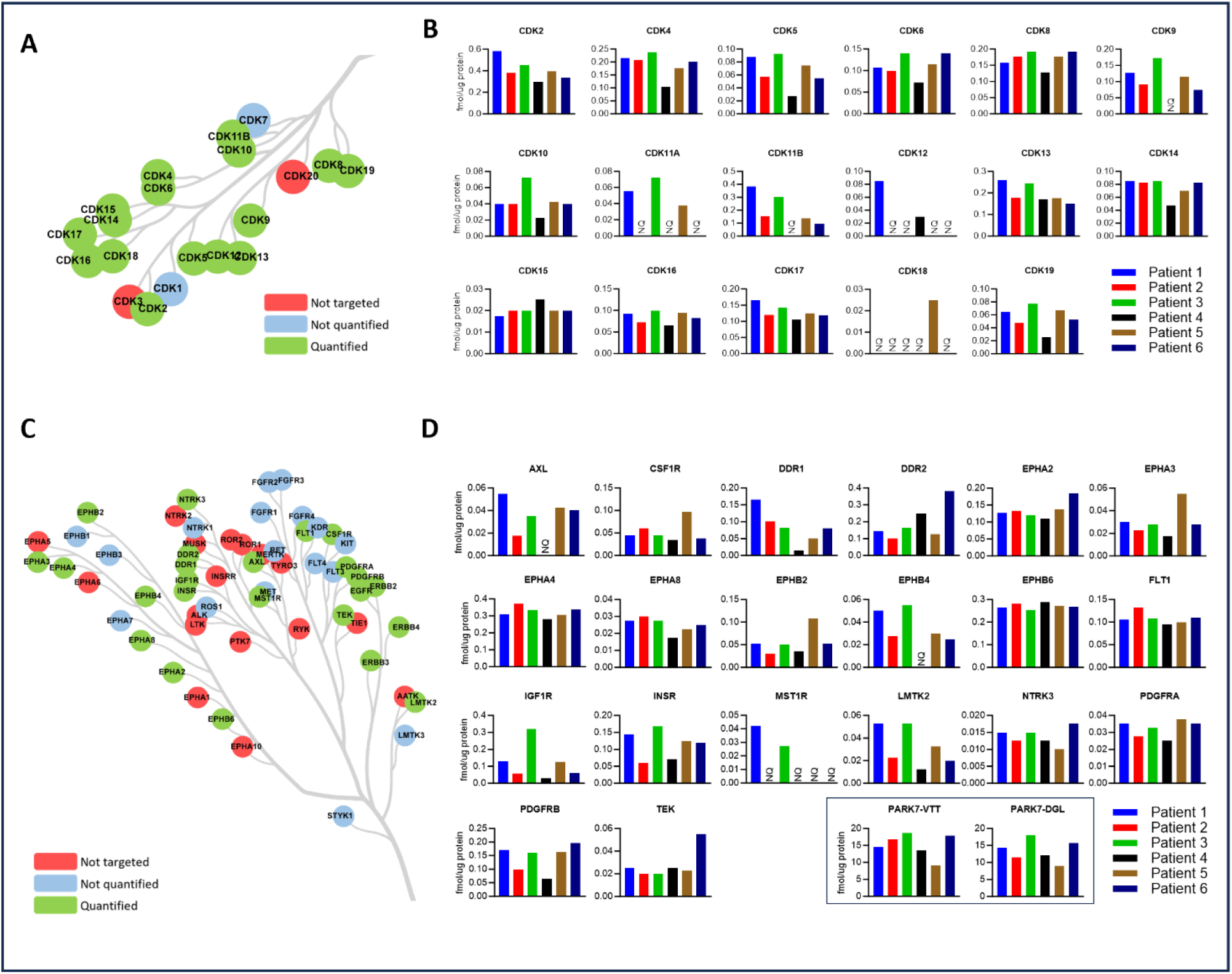
Quantitation of kinase protein expression within kinome sub-families by SureQuant/PRM in flash-frozen breast cancer patient tumors. **A)** The human genome encodes for 20 cyclin dependent kinases (CDKs). The current SIL library targets 18 CDKs of which 16 were quantified in at least one patient tumor. **B)** SureQuant/PRM quantitation of CDKs across patient tumors. **C)** The current SIL library targets 41 of the 58 receptor tyrosine kinases (RTKs) in humans. 24 RTKs were detected in at least one patient sample. **D)** SureQuant/PRM quantitation of RTK protein abundance in patient tumors. For both CDKs (A) and RTKs (C) green circles indicate proteins that were quantified in at least one sample, blue circles indicate proteins targeted by our SIL library but that were not quantifiable, and red circles indicate proteins for which we currently have no corresponding SIL. “NQ” indicates that a protein could not be quantified in the indicated sample due to low abundance, interference by unrelated product ions, or an insufficient number of points across product ion peaks. Quantitation of kinase proteins in (B) and (D) in each patient sample were from the same MS run. In the box at the bottom of (D) two peptides for PARK7/DJ-1 protein are shown as quantitative standards for control of total protein abundance for each patient tumor^37^.

RTKs respond to extracellular stimuli and perturbations to regulate intracellular signaling pathways. Many RTKs have selective expression profiles across different cell types and/or tissues and are frequently mutated or amplified in human cancers to drive cell proliferation and transformation. Our SIL library currently targets 41 of the 58 human RTKs, with SIL selection for the remaining 17 currently in progress (Fig. 2C). Of the 41 RTKs targeted, we quantified abundance of 24 across breast cancer patient samples (Fig. 2D). A subset of RTKs were expressed at similar levels across all patient breast cancer tumors including EPHA2, EPHA4, EPHB6, FLT1, NTRK3, and PDGFRA. Expression of the remaining RTKs was heterogeneous with most exhibiting ≥ 2-fold range in protein expression. Of the heterogenous RTKs, AXL, CSF1R, and IGF1R are therapeutic targets. DDR1, EPHA3, EPHA8, INSR, MST1R, PDGFRB, and TEK are each prognostic indicators for different cancers and DDR2 expression may enhance anti-PD1 immunotherapy^30-36^, suggesting quantitation of DDR2 protein expression is of significance for efficacy of immune checkpoint therapies. The deglycase protein PARK7 is expressed with low variability between cell types and at lower, more relevant levels than most other housekeeping genes like actin or GAPDH. Thus, PARK7 is perhaps the most reliable marker for cellular protein/peptide normalization in targeted proteomics experiments^37^. The box at the bottom of Figure 2D shows quantitation of two peptides for PARK7 demonstrating approximately equivalent total cellular protein/peptide abundance across patient samples (Suppl Table 2, Figs. 2B, D and 3A, B).

Figure 3A shows the clinical subtyping for ERBB2/HER2 of the same flash frozen patient tumors used for SureQuant/PRM in both Figures 2 and 3B. Of the six patients, 1 and 2 were ER+/PR+/HER2+, patients 3 and 4 were ER+/PR+/HER2-, patient 5 was ER+/PR-/HER2-, and patient 6 was ER-/PR-/HER2-. Clinically, ERBB2/HER2 expression status is currently defined by immunohistochemistry (IHC) staining of fixed tissue samples on microscope slides. Thus, assessment is not generally measured in comparison to other ERBB proteins or a proteomic normalization standard such as PARK7. The IHC scoring scale incorporates both abundance and subcellular localization ranging from 0-3+ with 3+ representing the highest level of expression and 0 representing no or barely detectable staining intensity^38^. Patients with ERBB2/HER2 scores of 3+, or 2+ with positive HER2 amplification on *in situ* DNA-based hybridization staining (FISH), are candidates for ERBB2/HER2 targeted therapies. Importantly, results from the Destiny-Breast04 Clinical Trial established the antibody drug conjugate Trastuzumab Deruxtecan (T-DXd) as a new standard of care for HER2-low metastatic breast cancer, where ERBB2/HER2-low is a new standard representing 1+ and 2+/FISH-negative patients based upon the classical ERBB2/HER2 clinical assay. ER/PR negative^39^, HER2-low patients that were categorized as Triple Negative Breast Cancer (TNBC) responded to T-DXd with nearly a 3-fold longer progression free survival (8.5 months versus 2.9 months) and > 2-fold longer overall survival (18.2 months versus 8.3 months) compared to chemotherapy alone. Within this group, the percentage of patients with a confirmed objective response to T-DXd was 50%. Thus, a quantitative measure of kinase protein (ERBB2/HER2) abundance in patients’ tumors would significantly augment therapeutic decisions based on IHC.

**Figure 3:**
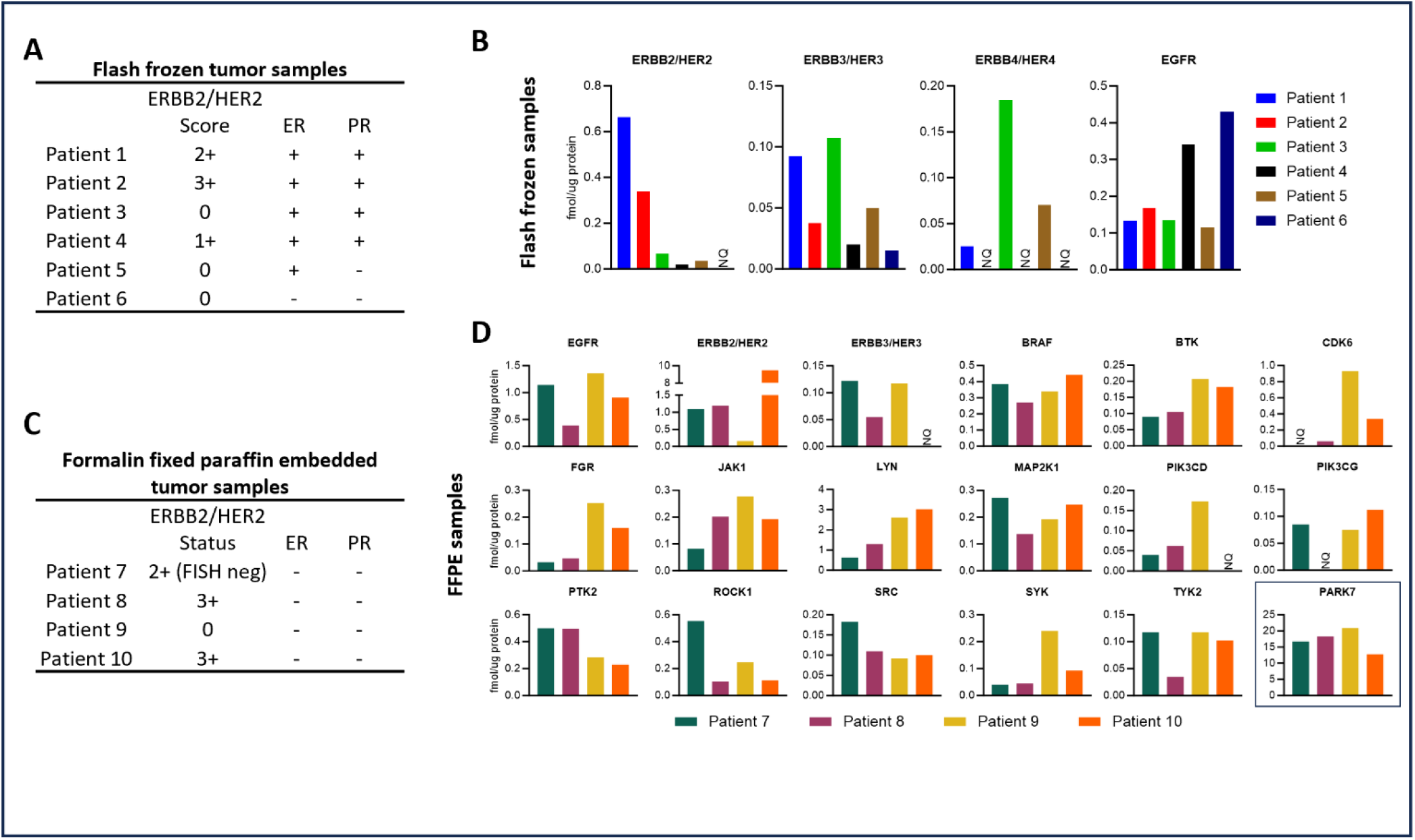
Quantitation of kinases relevant to oncogenesis and targeted pharmacological inhibition from flash frozen and formalin fixed paraffin embedded breast cancer patient tumors. **A)** Clinical determination of estrogen receptor (ER) and progesterone receptor (PR) status and ERBB2/HER2 scoring by immunohistochemistry (IHC) of flash frozen tumor samples. **B)** Quantitation of the EGFR/ERBB family of receptor tyrosine kinases by SureQuant/PRM in flash frozen tumor tissue samples described in (A). Patient tumor sample analysis in panels A and B for the EGFR/ERBB kinases are from the same MS run as the CDK and RTK analysis in Figure 2 including the PARK7 controls and can be directly compared for expression profiles. **C)** Clinical determination of ER, PR, and ERBB2/HER2 status by IHC in formalin fixed paraffin embedded (FFPE) tumor samples. **D)** Quantitation of clinically relevant kinases quantified by SureQuant/PRM from FFPE tumor samples described in (C). “NQ” indicates that a protein could not be quantified in the indicated sample due to low abundance, interference by unrelated product ions, or an insufficient number of points across product ion peaks. Quantitation of kinase proteins in (D) were from the same MS run for each patient sample.

In our SureQuant/PRM quantitation of the EGFR/ERBB RTK family (Figs. 3A and 3B), ERBB2/HER2 was quantified in five of the six patient tumor samples analyzed despite IHC scores of 0 for three patients. Patients 1, 2, and 4 had IHC scores of 2+, 3+, and 1+ respectively, indicating a wide range of ERBB2/HER2 expression across patients by IHC. Our SureQuant/PRM analysis showed a 33-fold range in ERBB2/HER2 protein levels but did not fully substantiate IHC scoring in HER2+ patients. Patient 4 had the lowest amount of detectable ERBB2/HER2 and we detected approximately 2-fold more ERBB2/HER2 in patient 1 than in patient 2 despite patient 2 having a higher IHC score. SureQuant/PRM shown in Figure 3B gives the quantitative measure of not only ERBB2/HER2 but also for the other ERBB family kinases including EGFR, ERBB3/HER3, the major dimerization partner for ERBB2/HER2, and ERBB4/HER4. Considering the relatively similar cellular protein/peptide loading based on PARK7 quantitative measurements (Fig. 2D), we saw significant heterogeneity of ERBB family RTK expression across the six breast cancer tumors.

We quantified kinase abundance from formalin fixed paraffin embedded (FFPE) tumor samples from four additional patients (Fig. 3C, D). Panel C shows patients 8 and 10 were ERBB2/HER2-high and patient 9 ERBB2/HER2 negative. Patient 7 was more equivocal, scoring 1+ with pretreatment biopsy and then 2+ but FISH negative in a second analysis. Consistent with flash frozen tumors, the FFPE tumors had a wide range (∼60-fold) in ERBB2/HER2 protein levels across samples including detection in samples that were scored as low by IHC. Of the two ERBB2/HER2-high patients by IHC, expression of ERBB2/HER2 in patient 10 was almost 10-fold greater than patient 8. Patient 7, which by IHC scored ERBB2/HER2-low, expressed similar levels of ERBB2/HER2 by SureQuant/PRM as patient 8 that was scored ERBB2/HER2-high. PARK7 quantitation in panel 3D shows these differences are not due to a significant discrepancy in proteomic normalization between samples prepared for LC-MS. SureQuant/PRM quantitation of other clinically relevant protein kinases targeted by small molecule inhibitors are also shown in Figure 3D and demonstrate heterogeneity of protein expression in FFPE patient tumor samples. Approximately 175 kinases and kinase-associated proteins per sample were quantified from FFPE slices (Suppl Table 3).

### Heterogeneity of protein kinase expression in Triple Negative Breast Cancer and ERBB2/HER2+ Subtypes

Clinical diagnosis of breast cancer subtypes is based on IHC scoring of ER, PR, and ERBB2/HER2 (see Fig. 3A), which guides treatment decisions. TNBC is generally characterized by the lack of ER and PR and varying levels of HER2 that may range from 2+ to undetectable levels of ERBB2/HER2 expression by IHC. Thus, TNBC is described as a very heterogenous disease^40^. Genomic analysis can categorize breast cancer into molecular subtypes which generally correspond with receptor subtypes^41^, but quantitative proteomic characterization of the heterogeneity of proteomic kinase expression is lacking. We have shown previously that subtypes of breast cancer can be defined by kinase taxonomies determined by MIB/MS functional kinase expression profiles ^10^, but quantitative measurements of kinase abundance were not possible with these methods. To demonstrate the quantitative measure of kinase protein expression, we used SureQuant/PRM to characterize kinase expression signatures of HER2 enriched (HER2e) and TNBC using cultured cell lines (Suppl table 4). Figures 4A and 4B show the quantitative measurement of differences in kinase expression in nine TNBC cell lines (HCC70, HCC1143, HCC1806, Hs578T, MDA-MB-231, MDA-MB-468, SUM102PT, SUM149PT, and SUM159PT) versus two HER2e cell lines (SKBR3, BT474). The volcano plot in Figure 4A highlights the kinases that were ≥2-fold enriched in TNBC (purple) versus HER2e (green) cells. As expected, ERBB2/HER2 was the most heavily enriched kinase in HER2e cell lines. The distribution of quantified expression levels across the cell lines are shown in Figure 4B. The size of the boxes for the different kinases indicates heterogeneity not just between TNBC and HER2e cells but within the cell lines of the two breast cancer subtypes with larger boxes indicating greater heterogeneity in kinase expression. We next examined heterogeneity in kinase abundance across two intrinsic subtypes of TNBC, basal-like (HCC70, HCC1143, HCC1806, MDA-MB-468, SUM102PT, and SUM149PT) and claudin-low (Hs578T, MDA-MB-231, and SUM159PT) (Fig. 4C, D). Kinases enriched in the basal-like TNBC subtype included the transcriptional regulator CDK19, three receptor tyrosine kinases DDR1, IGF1R, and MST1R, and one non-receptor tyrosine kinase TNK1. Kinases enriched in claudin-low cell lines were more varied but included two receptor tyrosine kinases AXL and EPHA2, the transcriptional regulator CDK7, and the NFκB pathway kinase IKBKB. Many of the kinases enriched in basal-like or claudin-low TNBC subtypes have been implicated in different human cancers.

**Figure 4:**
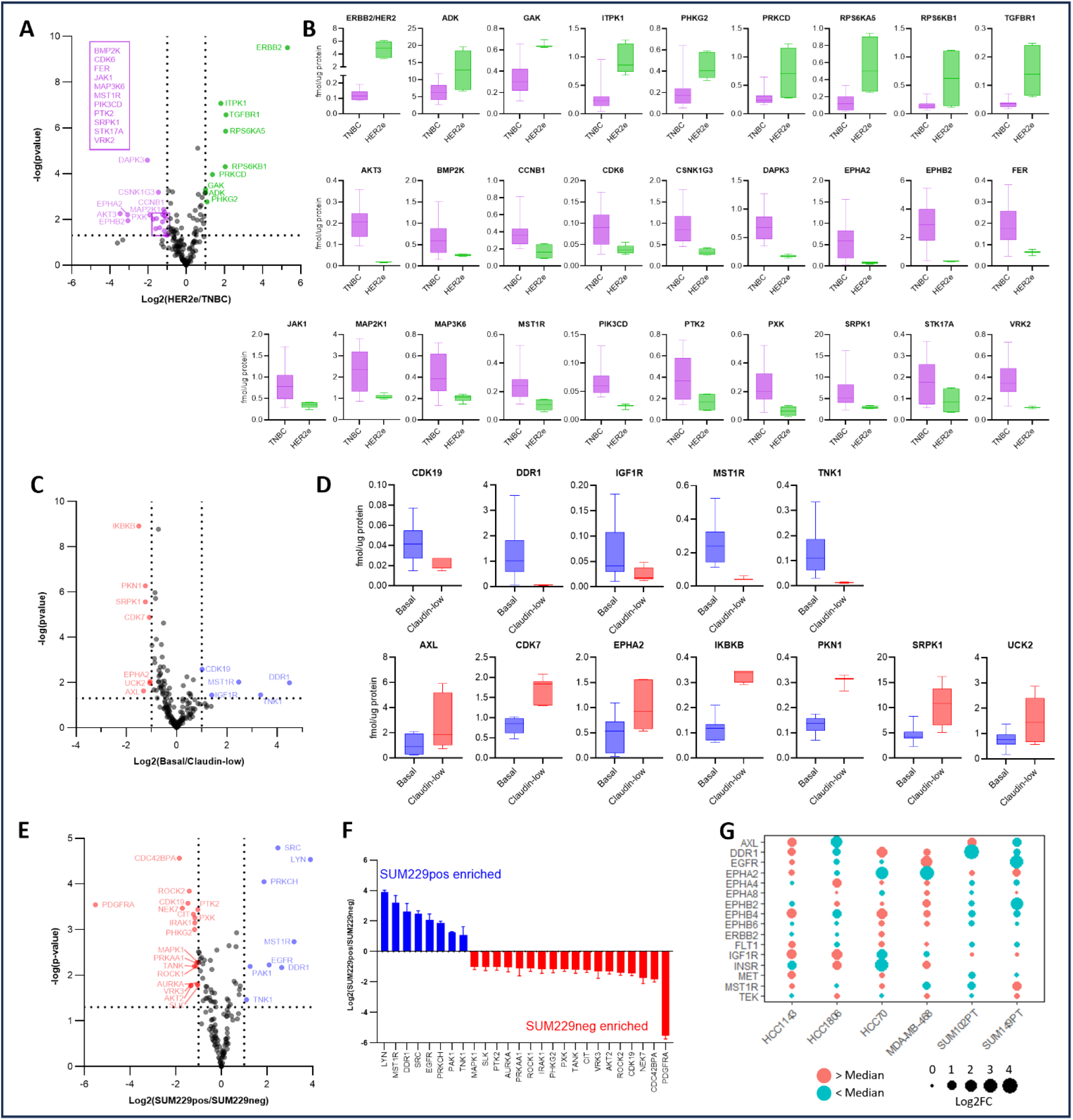
Differential expression of kinases across and within intrinsic subtypes of breast cancer cell lines. **A)** Panel of nine triple negative breast cancer (TNBC) and two HER2-enriched (HER2e) breast cancer cell lines were analyzed by SureQuant/PRM. Cell line breast cancer subtypes: TNBC basal-like: HCC70, HCC1143, HCC1806, MDA-MB-468, SUM102PT, SUM149PT; TNBC claudin-low: Hs578T, MDA-MB-231, SUM159PT; HER2e: SKBR3, BT474. The volcano plot indicates differential expression of kinases between TNBC and HER2e breast cancer subtypes. Kinases that were statistically significantly enriched by at least 2-fold in HER2e (green) or TNBC (purple) cell lines are highlighted. **B)** Quantitation of kinase protein expression of the differentially enriched kinases across HER2e and TNBC cell lines identified in (A). **C)** The 9 TNBC cell lines from (A) were subdivided by their basal (blue) and claudin-low (red) intrinsic molecular subtypes and kinase protein expression compared as in (A). **D)** Quantitation of kinase expression levels across basal and claudin-low cell lines of differentially expressed kinases identified in (C). **E)** The TNBC cell line SUM229PE consists of two phenotypic subpopulations that can be separated based on high (SUM229pos) or low (SUM229neg) expression of EPCAM and CD49f that can be cultured independently^43^. SUM229pos and SUM229neg cells were analyzed by SureQuant/PRM for differential kinase expression as in (A). **F)** Relative expression levels of differentially expressed kinases in each SUM229PE subpopulation. Three kinases enriched in basal TNBC cell lines (C and D), MST1R, DDR1, and TNK1 were also enriched in SUM229pos cells. **G)** Relative protein expression levels of receptor tyrosine kinases from the six basal TNBC cell lines characterized in panels A-D are shown in the dot plot. Circle size correlates with log2 transformed fold difference from the median expression level. Red circles indicate protein levels higher than the median while blue circles indicate protein levels lower than the median. All samples were analyzed in biological triplicate. Box and whisker plots (B and D) indicate the minimum and maximum values (error bars), 25^th^ – 75^th^ percentile (boxes), and median (line).

To emphasize further the heterogeneity of kinase expression in closely related tumor cell populations, we used the SUM229PT TNBC cell line that, with proper growth conditions *in vitro*, maintain two distinct phenotypic subpopulations^42,43^. One population has epithelial characteristics similar to the basal-like subtype (SUM229pos), and a second population with more mesenchymal characteristics similar to the claudin-low subtype (SUM229neg)^42,44^. SUM229pos and SUM229neg cells are genomically highly similar with differences in open chromatin states and enhancer-dependent transcriptional control regulating the phenotypic differences in the two populations^42^. Isolated populations of SUM229pos and SUM229neg cells were analyzed for quantitative kinase expression profiles using SureQuant/PRM and demonstrated considerable heterogeneity (Fig. 4E, F, Suppl table 5). Consistent with the basal-like characteristics, the three kinases (MST1R, DDR1, and TNK1) enriched in basal cell lines from Figures 4C and D were also enriched in SUM229pos cells.

Receptor tyrosine kinases are some of the most differentially expressed kinases across cell types and are implicated in a variety of cancers^45^. EGFR is overexpressed in ∼22% of TNBC and epigenetic up-regulation of other receptor tyrosine kinases is a major mechanism of adaptive resistance to targeted therapies^46,47^. We analyzed the proteomic expression heterogeneity in receptor tyrosine kinases within the six basal-like TNBC cell lines (Fig. 4G). EGFR had a nearly 100-fold difference in expression between the highest and lowest expressing cell lines. Heterogeneity of receptor tyrosine kinase protein expression within the basal-like cell lines rivaled the level of heterogeneity across breast cancer subtypes, consistent with what was observed in patient biopsies (Figs. 2 and 3), demonstrating why single targeted RTK inhibitors or combinations of targeted kinase inhibitors has been ineffective for treatment of TNBC.

### Integration of phospho-Ser/Thr proteomics with kinase SureQuant/PRM

We have identified more than 15,000 phosphosites using only 100μg protein from human TNBC organoids (Suppl Table 6). Fig. 5A and B show the heat maps for SureQuant/PRM analysis and phosphopetides for three different patient TNBC organoids (referred to as NH85TSc, NH87TT and NH95TT^48^). The organoids are established and free of stromal cells. The findings show differential patterns of kinases in each organoid line, consistent with the heterogeneity measured for human TNBC tumors and cell lines (Fig. 5A compared to Figs. 2-4, Suppl Table 7). The phosphoproteomes of the three organoid lines show similar heterogeneity with >8,000 differentially expressed phosphosites across the three organoid lines by ANOVA multiple-sample statistical analysis (Fig. 5B). Phosphorylation of a kinase may affect its enzymatic activity, localization, or association with binding partners or substrate proteins, all of which may contribute to its cellular functions. Thus, we integrated phosphoproteomics with quantitative measure of kinase abundance via SureQuant/PRM. We quantified phosphorylation state differences across organoids for a total of 240 kinases and quantified abundance of 206 kinases by SureQuant/PRM in parallel resulting in 121 kinases with matched phosphorylation state and SureQuant/PRM abundance (Fig. 5C). A total of 741 phosphorylation events on kinases were detected of which 402 were from kinases with matched SureQuant/PRM abundance (Fig. 5D). Of the over 90,000 phosphosites reported across the human proteome, only a small fraction have known biological consequences on their respective proteins. Using the phosphosite database from PhosphoSitePlus^49^, we annotated 189 phosphosites with documented regulatory functions including 115 from SureQuant/PRM quantified kinases (Fig. 5D). Pairwise comparisons across the three organoid lines mapping the relative abundance of kinase phosphosites against SureQuant/PRM quantitative abundance of kinases showed a strong correlation between kinase abundance and phosphorylation state (Fig. 5E-G). Phosphosites with regulatory function are highlighted in red and also demonstrate a strong correlation with kinase abundance suggesting that quantitation by SureQuant/PRM is representative of functional kinase expression. Quantitation of terminal kinases in two of the primary proliferative signaling pathways, MEK/ERK and AKT/mTOR suggested differential activation between the three organoid lines. RPS6KA1 and RPS6KA4 are members of the MEK/ERK signaling pathway and demonstrate higher levels of abundance and phosphorylation in the NH85TSc organoid line (Fig. 5E and F). RPS6KB1 and RPS6KB2 are in the AKT/mTOR signaling pathway and are significantly elevated in the NH95TT and NH87TT compared to the NH85TSc line.

**Figure 5:**
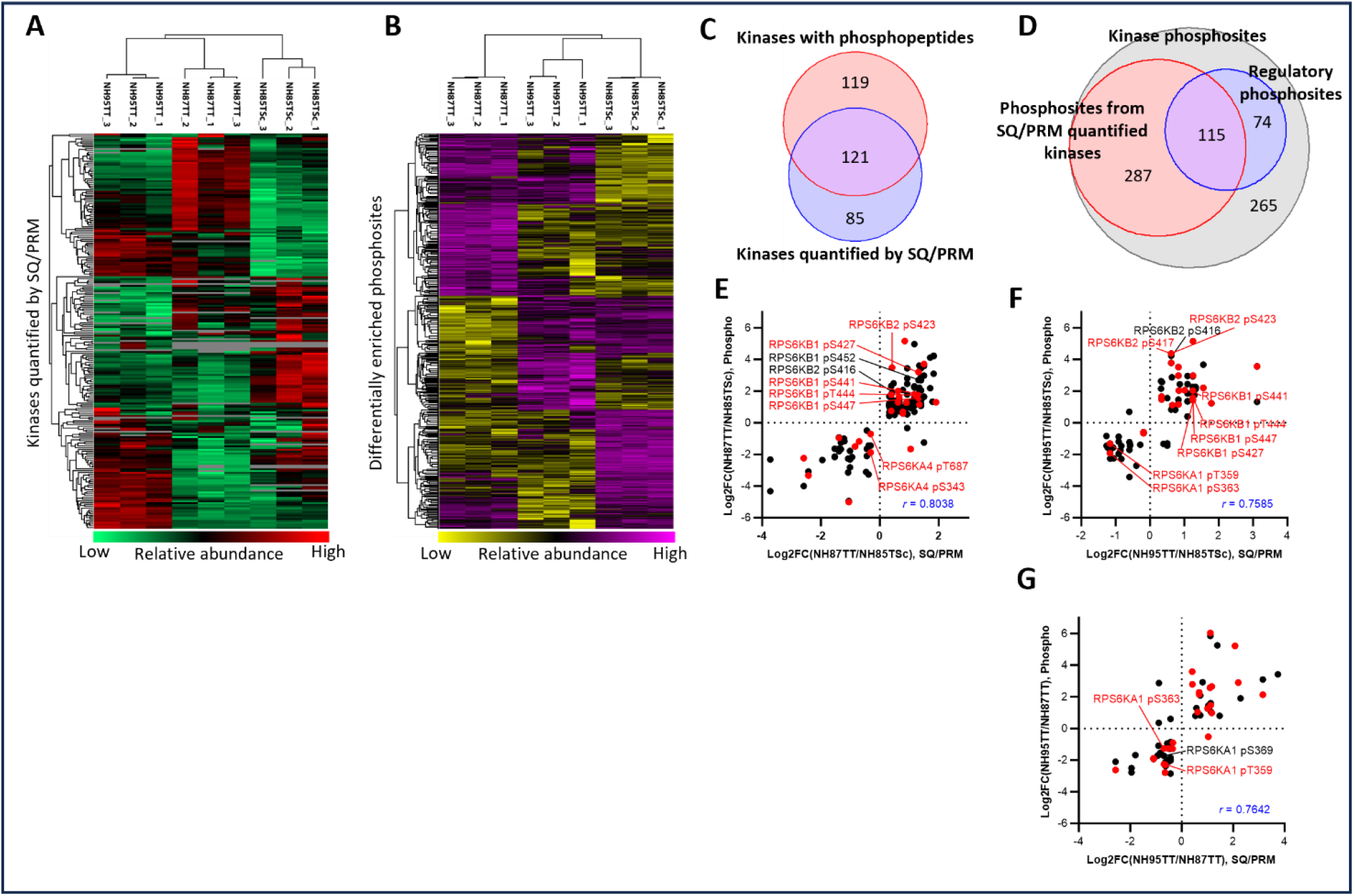
Phosphoproteomics reveals correlation between kinase abundance and phosphorylation in patient derived organoids. Three TNBC patient derived organoid (PDO) lines were analyzed by SureQuant/PRM and nanoscale phosphoproteomics^48,50^. **A)** The heat map of kinase protein expression demonstrates the heterogeneity of the kinome between the three PDO lines. **B)** Phosphoproteomics showed distinct patterns of protein phosphorylation across the three PDO lines. Shown are differentially enriched phosphosites determined by analysis of variance (ANOVA). **C)** Phosphosites were identified for 240 kinases of which 121 were quantified by SureQuant/PRM. **D)** Of the 741 phosphosites on kinases detected, 402 were from kinases quantified by SureQuant/PRM including 115 phosphosites known to have a regulatory effect on their respective kinase. **E-G**) Relative abundance of total kinase protein was plotted against the relative abundance of phosphorylation events on those kinases for each pairwise comparison between the three PDO lines. Only statistically significantly different phosphosites and kinase proteins are shown. Red circles indicate phosphosites with defined regulatory functions^49^

Tsai et al^50^ have developed tandem tip sample preparation methods for enrichment of phospho-Ser/Thr tryptic peptides that allows for nanoscale phosphoproteomic analysis. Using these nanoscale methods, it is possible to easily integrate kinase SureQuant/PRM with phospho-Ser/Thr proteomics (Extended Data Fig. 2). Using 10μg of starting organoid protein >9,000 phosphosites were identified across the three TNBC organoids. Filtering to retain only phosphosites quantified in all three biological replicates in at least one organoid line yielded > 6000 phosphosites for further analysis (Extended Data Fig. 2A, Suppl table 8). The specific phosphosites quantified were largely overlapping with the nanoscale phosphopeptide enrichment offering approximately half the coverage of total, regulatory, and kinase phosphosites (Extended Data Fig. 2A-C). Similar correlations between total protein abundance and phosphosite abundance were observed (Extended Data Fig. 2D-F) and overlaying coverage of phosphorylated proteins with the “Pathways in Cancer” KEGG pathway demonstrated nearly as much breadth of coverage (Extended Data Fig. 2G-H). Thus, as little as 20μg protein from the same starting tumor source allows an integrated SureQuant/PRM quantitative measure of expressed protein kinases and phosphoproteomics.

### Substrate predictions based on the quantitation of kinase expression using kinome SureQuant/PRM

“An atlas of substrate specificities for the human Ser/Thr kinome” was recently published by Johnson et al^51^, defining the substrate landscape of the human serine/threonine kinome based on primary structure preferences of purified kinases. We used this atlas of substrate specificities to predict upregulated phosphorylation of substrates based on the kinases specifically enriched in the NH95TT PDO line (Fig. 6A). Gene set enrichment analysis was performed on the predicted substrates using the Broad Institute’s Molecular Signatures Database of hallmark and curated gene sets^52^. A strong collective enrichment of gene sets associated with the immune response/system was observed (Fig. 6B). We hypothesized that either an immune cell population persisted within the NH95TT PDO line that was absent in the NH85TSc and NH87TT PDO populations or a viral or bacterial infection was present in the NH95TT PDO line. To test these hypotheses, we performed single nucleus RNAseq and ATACseq of the NH95TT and NH87TT PDO lines (Fig. 6C). Using the Kraken bioinformatics tool^53^ to detect alignment of RNA/DNAseq data with the genomes of > 8,500 archaeal, bacterial, and viral pathogens, we found no evidence of contamination/infection in the NH95TT PDO line (data not shown). The InferCNV computational tool (https://github.com/broadinstitute/inferCNV) was used to identify possible evidence of somatic chromosomal copy number alterations (CNAs) based on NH95TT single nucleus RNA sequencing. CNAs are much more frequent in tumor cells versus non-transformed cells, allowing for the computational prediction of “normal” and “tumor” cells from scRNAseq datasets. There was no evidence of subpopulations with fewer CNAs suggesting that all cells within NH95TT were tumor cells and no other “normal” cell populations existed (data not shown). Thus, the immune signatures observed from kinase substrate predictions were intrinsic features of NH95TT tumor cells.

**Figure 6:**
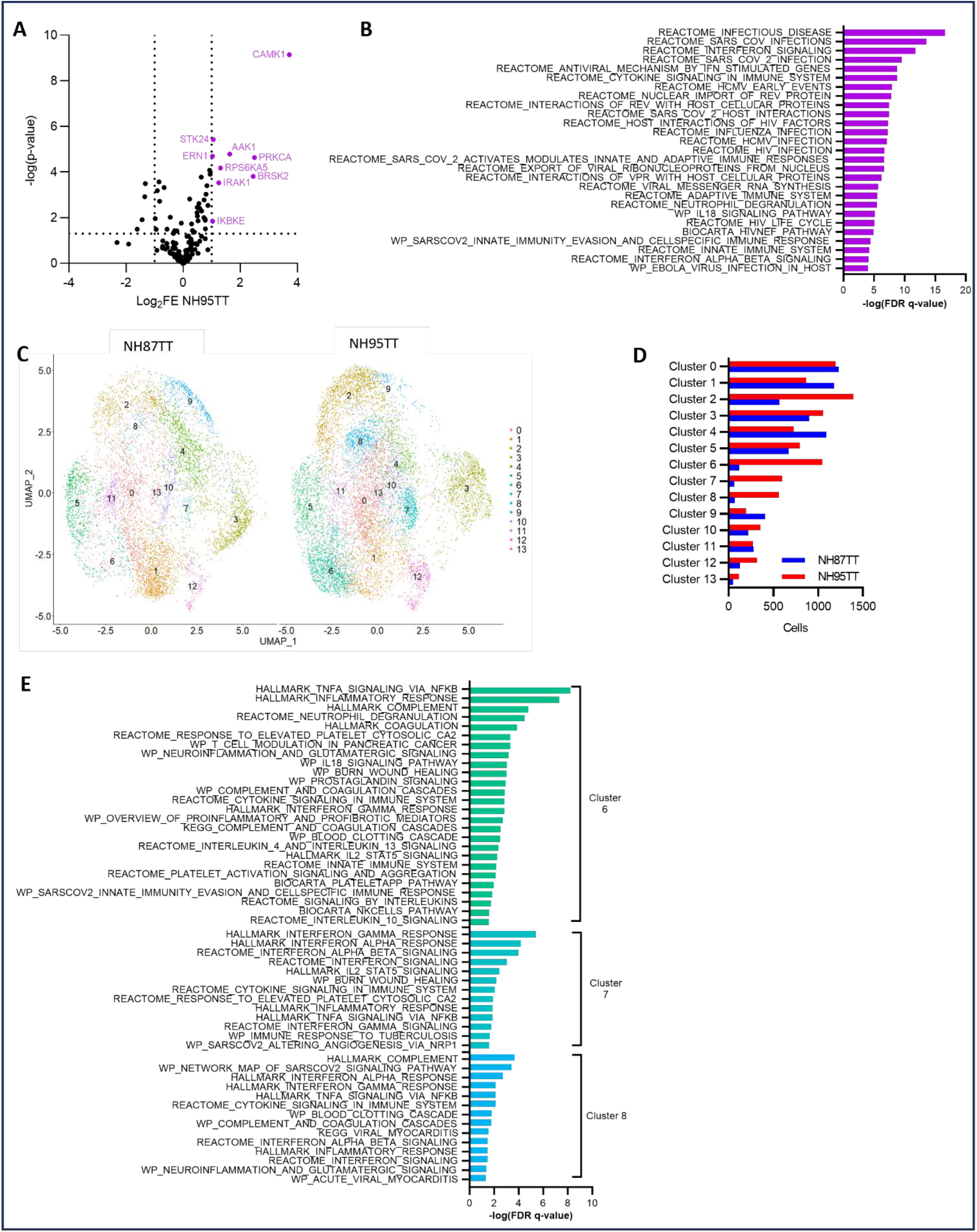
Kinase substrate predictions based on differentially expressed kinases determined phenotypic states in patient derived organoids. **A)** SureQuant/PRM analysis of three TNBC patient derived organoid (PDO) lines from Figure 5 showed enrichment of nine serine/threonine kinases in the NH95TT line when compared to the NH85TSc and NH87TT lines. **B)** The substrate specificity atlas for Ser/Thr kinases^51^ was used to predict putative substrates for enriched kinases from (A). Gene set enrichment analysis (GSEA) was performed on the putative kinase substrates using the Broad Institute’s molecular signature database^71^ demonstrating gene sets associated with immune and inflammatory signatures were heavily enriched in the NH95TT organoid line. **C)** Single nucleus RNA sequencing was performed on the NH95TT and NH87TT PDO lines and distinct cell clusters called using the Seurat R package^54^. Samples were integrated before clustering but plotted separately. **D)** Cell clusters 6, 7, and 8 from (C) were comprised almost exclusively of NH95TT cells. **E)** GSEA of genes enriched in clusters 6, 7, or 8 showed enrichment of immune and inflammatory associated gene sets as predicted in the proteome analysis of substrates shown in panel B.

The Seurat R toolkit^54^ was used to analyze single nucleus RNAseq of NH87TT and NH95TT PDO lines. Gene expression matrices were integrated to define conserved cell populations between the two PDO lines resulting in 14 distinct cell subpopulations (Fig. 6C). The distribution of cells in each cell cluster was approximately equivalent for each of the PDO lines (Fig. 6D) except for clusters 6, 7, and 8, which were heavily enriched for NH95TT cells. Genes enriched in each of the 6, 7, and 8 clusters were subjected to gene set enrichment analysis as in Figure 6B. In each of these clusters, we observed enrichment of gene sets associated with the immune system/response (Fig. 6E). These data are consistent with the immune signature predicted by the kinase substrate prediction analysis (Fig. 6B) and suggest that immune signatures derive from distinct cellular subpopulations of tumor cells within the NH95TT PDO line. Thus, the phenotype predictions from kinase substrate computational analysis based on SureQuant/PRM quantitation of the human kinome were substantiated by single nucleus RNAseq.

## Discussion

Here we describe a stable isotope labeled peptide library for internal standard PRM LC-MS coupled with SureQuant triggered acquisition software for quantitative measurement of >300 protein kinases in the human kinome. SureQuant/PRM gives an unbiased measurement of kinase expression at attomole-femtomole levels that requires no prior enrichment from flash frozen or FFPE tissues. Ten μg or less tissue protein is required for LC-MS analysis of tryptic peptides, allowing quantitative measurements of clinically relevant kinase expression from patient biopsies. Kinome Surequant/PRM studies in breast cancer provided a precise quantitative measure of the heterogeneity in kinase protein expression in patient biopsies. Heterogeneity was similarly observed comparing kinase expression levels within HER2-enriched and TNBC cell lines and organoids.

The clinical relevance of SureQuant/PRM proteomic analysis of protein kinases was clearly shown with TNBC patient tumor biopsies (Figs. 2 and 3). It was possible to quantitate clinically relevant families of kinases such as CDKs and both tyrosine and serine/threonine kinases that have FDA approved inhibitors for treatment of different cancers in both flash-frozen needle biopsies and FFPE tissue samples. Of significant clinical relevance is the quantitation of the four ERBB family kinases (EGFR, ERBB2/HER2, ERBB3/HER3, ERBB4/HER4) where HER2 expression is measured in an attomole-femtomole scale rather than an IHC scale for HER2^38^. Quantitative measures of HER2 expression have become extremely important with the clinical findings that 50% of TNBC patients, considered HER2 low by IHC, respond to the ERBB2/HER2 targeting Trastuzumab-Deruxtecan antibody drug conjugate^39^. Using IHC analysis, it is presently unclear what low level of HER2 expression is required for a positive TNBC tumor response with Trastuzumab-Deruxtecan. It is also unclear if other members of the ERBB kinases, such as HER3 the preferred HER2 dimerization partner^55^, have a threshold of expression for functional response to Trastuzumab-Deruxtecan. Approximately 22% of TNBC tumors overexpress EGFR and having a quantitative measure of the four ERBB family members as well as an additional twenty or more RTKs, allows for a comprehensive understanding of the RTK landscape of each patient’s tumor. SureQuant/PRM of patient biopsies before and during treatment can address this important clinical question in the context of the human kinome.

In organoids, the ability to perform SureQuant/PRM and phospho-Ser/Thr proteomics from the same limiting sample allowed more than 15,000 phosphosites to be identified. Sixty kinases of the 226 quantified by SureQuant/PRM had phosphorylated regulatory sites identified that control different kinase functions (Fig. 5). The integrated datasets demonstrated kinase abundance correlated with the relative phosphorylation of the kinase, consistent with a strong relationship between kinase abundance and the regulation of signaling networks. We further validated this concept by using the “Atlas for substrate specificities for the human serine/threonine kinome”^51^ to predict pathways or phenotypic states that were differentially regulated in PDOs based on kinase expression heterogeneity (Fig. 6). This analysis indicated an enrichment in immune/inflammatory signatures in the NH95TT PDO that was substantiated by single nucleus RNA sequencing. Thus, comprehensive quantitative measurement of protein abundance of the kinome with SureQuant/PRM allows for determination of unique regulatory functions involved in the control of cellular physiology and disease states in addition to identifying putative targets for design of personalized treatment strategies.

In summary, SureQuant/PRM kinase analysis has the quantitative sensitivity to be used in the clinic to determine kinome expression characteristics in a needle biopsy or slice from an FFPE tumor. This novel capability permits the serial analysis of tumor samples to monitor cancer progression or response to treatment as well as comparison of metastases and primary tumor. Recent studies have emphasized the “proteome as a missing link between the genotype of oncogenic drivers and their functional state”^56^. Using needle biopsies for inhibitor pulldowns for kinase enrichment profiling^57^ and analysis of phosphorylated substrates^58,59^ clearly show the ability to integrate functional proteomics for clinical studies. As the proteomic datasets continue to expand and be refined, it will be possible to use quantitative SureQuant/PRM in the clinic with computational modeling to predict individual patient response to therapies^51,60-69^.

## Methods

### Cell culture

Cell lines and organoids were maintained at 37°C, 5% CO_2_. HCC1143, HCC1806, HCC70, Hs578T, SK-BR-3, and BT-474 were grown in RPMI supplemented with 10% fetal bovine serum (FBS) and 1% PennStrep (Gibco). MDA-MB-468 cells were grown in DMEM supplemented with 10% FBS and 1 % PennStrep. SUM102PT and SUM149PT cells were grown in complete HUMEC media supplemented with 10% FBS and 1% PennStrep. SUM159PT and MDA-MB-231 cells were grown in DMEM/F12 supplemented with 5% FBS, 1% PennStrep, 5 ug/mL insulin, and 1 ug/mL hydrocortisone. SUM229pos and SUM229neg cells were grown in F12 supplemented with 10 mM HEPES, 5% FBS, 5 ug/mL insulin, 1 ug/mL hydrocortisone, and 1% PennStrep. PDO lines were grown in 300 uL domes of Cultrex Ultimatrix reduced growth factor basement membrane (Bio-Techne), and maintained in DMEM/F12 (Gibco), 10% R-Spondin1 conditioned media obtained from Cultrex® HA-R-Spondin1-Fc 293T Cells (Bio-Techne), B27 supplement (Gibco), 5 mM Nicotinamide (Sigma-Aldrich), 1.25 mM N-Acetylcysteine (Sigma-Aldrich), 50 ug/ml Primocin (InvivoGen), 100 ng/ml Noggin (PEPROTECH), 5 ng/ml hEGF (PEPROTECH), 37.5 ng/ul hHeregulin-beta (PEPROTECH) 5 ng/ml FGF-7 (PEPROTECH), 20 ng/ml FGF10 (PEPROTECH), 5uM Y-27632 (Abmole Bioscience), 500 nM SB202190 (Sigma-Aldrich), 500 nM A83-01 (Tocris) as previously described^48^.

### Tumor samples

Patient tumor samples were provided by the UNC Tissue Procurement Core Facility as either flash-frozen needle biopsies or surgical resections of 10 um FFPE curls. All samples were collected with informed written consent from patients pursuant with the University of North Carolina’s Institutional Review Board.

### Stable isotope labeled peptides

SIL peptides were purchased from New England Peptides’ custom synthesis service at >95% chemical purity and 99% isotopic purity with uniformly C^13^ and N^15^ labeled C-terminal amino acids. Unlabeled peptides were also purchased at the same purity. Peptides were dissolved in 30% acetonitrile with 1% formic acid and combined at equimolar ratios then lyophilized and stored at -80°C until used. For the SureQuant survey run and all biological samples, SIL peptides were dissolved in 2% acetonitrile and 0.1% formic acid to a final concentration of 25 fmol/uL. For calibration curves, SIL peptides were reconstituted to a final concentration of 25 fmol/uL with varying concentrations of unlabeled peptides: 10 amol/uL, 25 amol/uL, 50 amol/uL, 125 amol/uL, 500 amol/uL, 1 fmol/uL, 5 fmol/uL, 25 fmol/uL, 50 fmol/uL, or 250 fmol/uL in 2% acetonitrile and 0.1% formic acid.

### Sample preparation

For SureQuant/PRM analysis, all cell culture or flash-frozen samples were lysed in buffer consisting of 50 mM Tris pH 8.0, 8 M urea, 75 mM NaCl, 1 mM EDTA, 2 ug/mL aprotinin, 10 ug/mL leupeptin, 1 mM phenylmethylsufonyl fluoride, 10 mM NaF, and 1% each of Sigma phosphatase inhibitor cocktails 2 and 3. Tumor samples were disrupted mechanically in lysis buffer with a motor driven pestle directly in microfuge tubes. Samples were sonicated for a total of 30 s in three 10 s bursts then clarified by centrifugation at 16,000 × g for 30 minutes at 8°C. Supernatants were collected in fresh microfuge tubes and protein concentration determined using bicinchoninic acid (BCA) assay (Pierce). 5 ug of total protein was reduced with 5 mM DTT at room temperature for 45 minutes then alkylated with 10 mM iodoacetamide in the dark for 45 minutes at room temperature. Samples were diluted 5-fold using 50 mM Tris pH 8.0 then digested with 0.1 ug Lys-C at room temperature for 2 hours. 0.1 ug of trypsin (Promega) was added and samples incubated overnight at room temperature. Proteolysis was quenched by acidification to pH < 2.0 with trifluoroacetic acid (TFA). 125 fmol (25 fmol/ug protein) of SIL library was spiked into each sample and samples desalted by solid phase extraction using Pierce peptide desalting columns according to manufacturer’s protocol. Eluted peptides were dried in speed-vac and stored at -80°C until LC-MS/MS analysis.

For all FFPE tumors, samples were deparaffinized by three 5-minute incubations with Xylene and rehydrated using three sequential washes of 100%, 85%, and 70% ethanol. The final pellet was resuspended in 100 mM ammonium bicarbonate, pH 8.0 and crosslinking reversed by incubation for two hours at 80°C. Samples were cooled to ambient temperature and an equal volume of 2,2,2 Trifluoroethanol (Sigma-Aldrich) was added. Samples were sonicated by five pulses of five seconds. Samples were incubated at 60°C for 60 minutes followed by another round of sonication. Protein concentration was determined using the BCA assay. Samples were then reduced in a final concentration of 13.3 mM Tris(2-carboxyethyl)phosphine (Sigma-Aldrich) and 33.3 mM DTT (Sigma-Aldrich) at 60°C for 30 minutes. Samples were then alkylated in 50 mM Iodoacetamide (Sigma-Aldrich) in the dark for 20 minutes at room temperature. Samples were diluted 2.5-fold with 50 mM ammonium bicarbonate, pH 8.0 and sequence grade modified trypsin (Promega) added at a ratio of 1:50 (w:w) to digest protein. Enzymatic digestion was quenched by acidification to pH < 2.0 with TFA and samples desalted using Pierce peptide desalting columns as per the manufacturer’s protocol. Samples were reconstituted in 2% acetonitrile and 0.1% formic acid and peptide concentration determined using the Pierce quantitative fluorometric peptide assay. SIL library was spiked in at a final concentration of 25 fmol per microgram of peptide and peptides dried in a speed-vac and stored at - 80°C until LC-MS/MS analysis^76^.

For phosphoproteomics, tryptic digests from PDO cell pellets were made using S-Trap micro spin column according to the manufacturer’s protocol (Protifi) except that TFA was used in place of phosphoric acid and lyophilized ^77^. Phosphopeptides were enriched using immobilized metal affinity chromatography as previously described ^50,78,79^. Peptides were reconstituted in 50% acetonitrile and 0.1% TFA until fully dissolved then acetonitrile with 0.1% TFA added to a final concentration of 80% acetonitrile. C18 stage tips were constructed using Empore C18 extraction disks and sequentially washed with methanol, then 50% acetonitrile with 0.1% formic acid. Stage tips were then conditioned with1% formic acid or 0.1% formic acid for the full-scale or nano-scale phosphopeptide enrichment respectively. Ni-NTA Superflow Agarose Beads were stripped of Ni with 100mM EDTA and replaced by incubation with FeCl_3_. Fe-NTA beads were washed and resuspended in an equal volume of a 1:1:1 solution of acetonitrile, methanol, and 0.01 % acetic acid for full-scale enrichment or with 1% acetic acid for nano-scale. 100 ug or 10 µg of tryptic peptide was mixed with 10 uL of Fe-NTA slurry and incubated at room temperature for 30 minutes with gentle mixing. Beads were spun at 1000 × g for 1 minute and supernatant discarded. Fe-NTA beads with enriched phosphopeptides were resuspended in 200 uL of 80% acetonitrile with 0.1% TFA and loaded onto the conditioned C18 stage tips. Stage tips were washed with 80% acetonitrile and 0.1% TFA then with 1% formic acid or 1% acetic acid for full-scale or nano-scale, respectively. Phosphopeptides were then eluted from the Fe-NTA beads onto the C18 plugs with withe 500 mM potassium phosphate pH 7.0 (full-scale) or with 200 mM NH_4_H_2_PO_4_ pH 4.4 (nano-scale) and C18 plugs washed with 1% formic acid. Phosphopeptides were eluted with 50% acetonitrile and 0.1% formic acid, dried in a speed-vac, and stored at -80°C until LC-MS/MS analysis.

### LC-MS/MS analysis

Samples were analyzed on a Thermo Orbitrap Exploris 480 mass spectrometer equipped with a Nanospray Flex ion source (Thermo) coupled with an UltiMate 3000 HPLC. Samples were injected onto a 15 cm Aurora Elite column (IonOpticks) heated to 40°C using a Sonation column oven. Buffer A consisted of 0.1% formic acid in water and Buffer B of 80% acetonitrile and 0.1% formic acid. Unless otherwise indicated, peptides were eluted over a linear gradient from 3-19% B over 72 minutes, 19-29% B over 28 minutes, 29-41% B over 20 minutes, 41-95% B over 3 minutes, and constant 95% B for 7 minutes at a constant flow rate of 250 nL/min. MS settings common to all runs included 2.1 kV spray voltage and 275°C ion transfer tube temperature.

An alternative gradient was used for all phosphoproteomics samples: 2 – 20% B over 100 minutes, 20 – 32% B over 20 minutes, 32 – 95% B over 1 minute, and constant 95% B for 19 minutes. Data dependent acquisition mode was used with 350 – 1800 scan range, 120,000 resolution, 100% normalized AGC target, and 60 ms injection time. For the top 15 ions, MS2 scan settings were 110 first mass (m/z), 15,000 resolution, 100% normalized AGC target, 105 ms inject time, and 30% higher energy collisional dissociation (HCD) collision energy.

### SureQuant/PRM LC-MS/MS

SIL library was spiked into a commercial HeLa tryptic digest (Thermo) at a concentration of 25 fmol/ug peptide, dried in a speed-vac, and reconstituted for a SureQuant survey run to determine optimal charge states, transitions, and triggering thresholds. A “Targeted Mass” list was provided including the z and m/z each peptide for each of the +2, +3, and +4 potential charge states. Full scan MS settings were set at 300 – 1500 m/z, 60,000 resolution, 300% normalized AGC target with a 40 ms inject time. Cycle time for full scan was set to 3 s with +/-10 ppm mass tolerance. MS2 scan settings included 150 – 1700 scan range, 7,500 resolution, 300% normalized AGC target, inject time of 40 ms and 30% HCD collision energy. Data were analyzed in Skyline and optimal charge states of SILs were defined as those with the greatest precursor intensities and triggering thresholds were set at 1% of the maximum peak height ^26,27^. The six most abundant product ions were selected for each precursor.

SureQuant quantitation methods were designed based on the information obtained for each peptide in the survey run and required detection of at least three of the specified product ions. The method consisted of 10 branches per full scan corresponding to a specific charge state of the specific heavily labeled amino acid and the optimal charge state from the survey run: A+2/3, E+3, K+2/3/4, R+2/3/4, and Y+2. Optimal charge state and corresponding m/z and intensity thresholds were provided for each precursor ion as “Targeted Mass” and their corresponding product ion m/z in “Targeted Mass Trigger.” The m/z ratios for up to six product ions were provided for “Targeted Mass Trigger.” Full scan MS settings included 300 – 1500 scan range, 120,000 resolution, 300% normalized AGC target, and 50 ms inject time. Data dependent acquisition was limited to 3 s cycle times with +/-3ppm mass tolerance. When a matching precursor ion was detected during MS1, a low resolution (7,500) MS2 scan of the SIL was triggered with 150 – 1700 scan range, 1000% normalized AGC target, inject time of 10 ms, and 27% HCD collision energy. Upon pseudomatching at least 3 product ions, a high resolution MS2 scan for the endogenous protein with relevant m/z offsets based on the heavy label was triggered with 150 – 1700 scan range, 60,000 resolution, 1000% normalized AGC target, 90 ms inject time, and 27% HCD collision energy.

### Mass spectrometry data analysis

All SureQuant/PRM data was analyzed using the Skyline software package ^26,27^. Peak area ratios were calculated for isotopically labeled heavy peptides versus the endogenous peptides for the three most abundant product ions with at least six points across the peak. Only two product ions were used to calculate peak area ratios in some cases of interference with a product ion. Peak area ratios were multiplied by the amount of SIL spike-in to determine absolute abundance levels. For all measurements, thresholds were applied for lower limit of quantitation thresholds based on calibration curves and 30% coefficient of variation across triplicate biological replicates applicable. Quantitation values failing to meet either threshold were discarded. Kinome trees were generated using the CORAL package^72^.

Phosphoproteomics data were analyzed using MaxQuant version 1.6.2.10 and the integrated Andromeda search engine using the SwissProt human proteome downloaded from Uniprot on 09/28/2024. Default MaxQuant settings were used with match between runs with a match time window of 4 minutes and an alignment window of 20 minutes. Variable modifications included oxidation (M), acetyl (protein N-term), and phospho STY. Fixed modications included carbamidomethyl (C). Only phosphosites with a localization probability >/= 0.75 and present in all three biological replicates in at least one sample group were considered. Phosphosite intensities were imported into the Perseus version 2.0.11 companion software and missing values imputed with a 0.3 width and 1.8 down shift. Data were normalized based on column median prior to statistical analysis for differential enrichment with ANOVA using a 0.05 FDR or with two-tailed students T-tests using *p* < 0.05 for pairwise analyses. Regulatory phosphosites were annotated based on the “Regulatory_sites” database downloaded from PhosphoSitePlus^49^.

### Kinase substrate predictions and gene set enrichment analysis

Supplementary Table 3 from the “Atlas for substrate specificities for the human serine/threonine kinome” was used for kinase substrate predictions^51^. Only substrates with a promiscuity score < 31, percentile ≥ 99, and a rank of 1 were considered. Gene set enrichment analysis was performed using the online portal for the molecular signatures database (https://www.gsea-msigdb.org/gsea/msigdb/index.jsp). Overlaps between enriched genes were computed against the hallmark and canonical pathways gene set collections.

### Single nucleus RNAseq and ATACseq

Single-cell suspensions from patient-derived organoid lines were made by digesting Cultrex domes with TrypLE, washing with PBS + 0.04% BSA, and filtering through Flowmi 40 µm cell strainers. Cells were spun at 300 × g for 5 minutes at 4°C, then resuspended by pipette mixing in chilled lysis buffer (10 mM Tris pH 7.4, 10 mM NaCl, 3 mM MgCl2, 0.1% Tween-20, 0.1% IGEPAL, 0.01% digitonin, 1% bovine serum albumin (BSA), 1 mM DTT, 1 U/uL RNase inhibitor) and incubated on ice for 5 minutes. Chilled wash buffer (10 mM Tris pH 7.4, 10 mM NaCl, 3 mM MgCl2, 0.1% Tween-20, 1% BSA, 1 mM DTT, 1 U/uL RNase inhibitor) was added to the samples, pipette mixed, and spun at 500 × g for 5 minutes at 4°C. The supernatant was discarded, and the pellet was washed twice with chilled wash buffer. The final nuclei pellet was resuspended in 1X Nuclei Buffer (10X Genomics) and immediately processed for library preparation using the 10X Genomics Chromium with the Single Cell Multiome ATAC + Gene Expression kit per the manufacturer’s protocol.

The resulting libraries were sequenced on an Illumina NextSeq 2000 instrument. Following sequencing, single nucleus RNAseq and ATACseq data were processed using the CellRanger software for demultiplexing, barcode processing and read counting. Single cell RNAseq and ATACseq data were further analyzed using the Seurat R package ^80^, inferCNV (https://github.com/broadinstitute/inferCNV), and/or the Kraken software package^81^.

The feature barcode matrices from CellRanger were converted to an R objects in Seurat. Nuclei were filtered to keep only those with UMI ≥ 500, genes ≥ 750, log(Genes/UMI) > 0.75, and < 20% mitochondrial counts. The matrix was further filtered to remove any genes that were detected in fewer than 10 cells. Count data were normalized, cell cycle status scored and regressed, and data scaled using Seurat’s NormalizeData, CellCycleScoring, and SCTransform functions. Data matrices were then integrated based on the most variable 3000 genes using SelectIntegrationFeatures, PrepSCTIntegration, FindIntegrationAnchors, and IntegrateData sequentially. Principle component analysis was run using RunPCA and cell clusters called using the first 40 principle components with FindNeighbers and FindClusters using a resolution of 0.6. Differentially expressed genes were determined using the FindAllMarkers function with a Log_2_FC threshold of 0.25.

## Acknowledgements

This work was supported by the UNC Lineberger Triple Negative Breast Cancer (TNBC) Center (GLJ), NIH U24DK116204 as part of the Illuminating the Druggable Genome Project (GLJ, RRT, SMG), NCI Breast SPORE P50CA058223 (CMP, HSE, LAC), NCI U01CA238475 (CMP, GLJ), NCI K08CA280388 (PMS), NCI R01CA233811 (SMG), NCI R01CA288145 (JJY, GLJ), NCI U01CA274298 (JJY, GLJ), Pancreatic SPORE P50CA25791 (JJY, GLJ), UNC LCCC Support Grant NCI P30CA016086 (HSE), UNC Lineberger Developmental Funding Program (MPE, DOO), NCI P01CA013106-Project 3 and CSHL/Northwell Health (DLS). We thank the UNC LCCC Tissue Procurement Core for breast cancer tumor specimens, the Translational Genomics Lab core facility (SCR_025231), and the UNC Metabolomics and Proteomics (MAP) Core for support of the Exploris 480 MS. Genomics project management was performed by the Lineberger Office of Genomics Research at the University of North Carolina-Chapel Hill, which is supported in part by the Lineberger University Cancer Research Fund, by the NIH NCI 5UG1CA233333 grant, and The UNC Center for Environmental Health and Susceptibility (UNC-CEHS) P30ES010126 grant. The Washington University Proteomics Shared Resource (WU-PSR) is supported in part by the WU Institute of Clinical and Translational Sciences (NCATS UL1TR000448), the Mass Spectrometry Research Resource (NIGMS P41GM103422) and the Siteman Comprehensive Cancer Center Support Grant (NCI P30CA091842) and CPTAC (U24CA160035). The expert technical assistance of Alan Davis and Rose Connors is gratefully acknowledged. Graphical abstract created with BioRender.com.

## Competing Interests

H.S.E. is a founder of Meryx (a UNC start-up) that is developing small molecule inhibitors for MERTK and owns stock in Meryx. C.M.P.is an equity stockholder and consultant of BioClassifier LLC; C.M.P. is also listed as an inventor on patent applications for the Breast PAM50 Subtyping assay. All other authors declare no conflicts of interest.

---

**Extended Data Figure 1:**
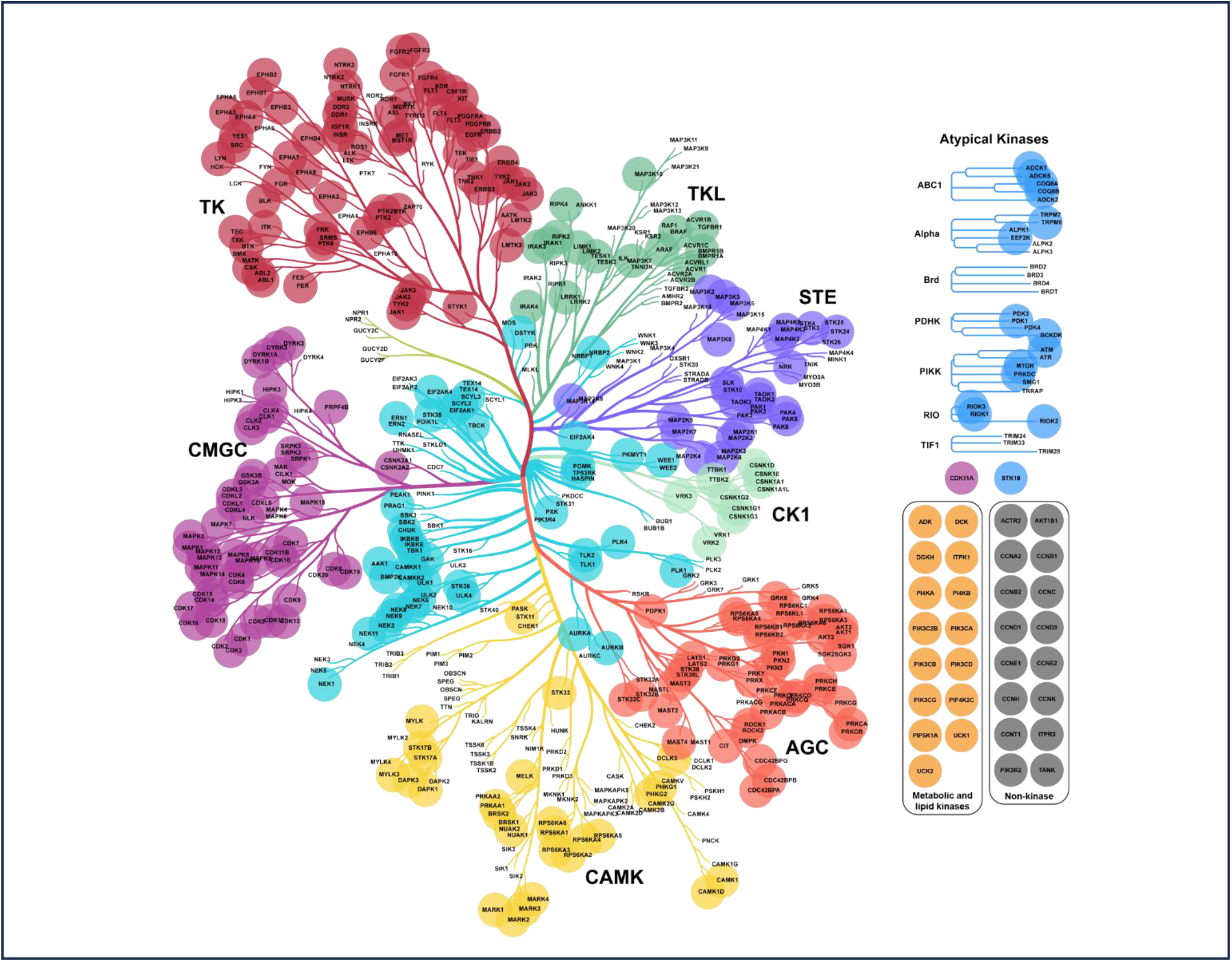
Kinome tree of SIL peptide library coverage. The kinases and kinase associated proteins targeted by our SIL peptide library are indicated by circles whose colors correspond to the kinase family^1,72^.

**Extended Data Figure 2:**
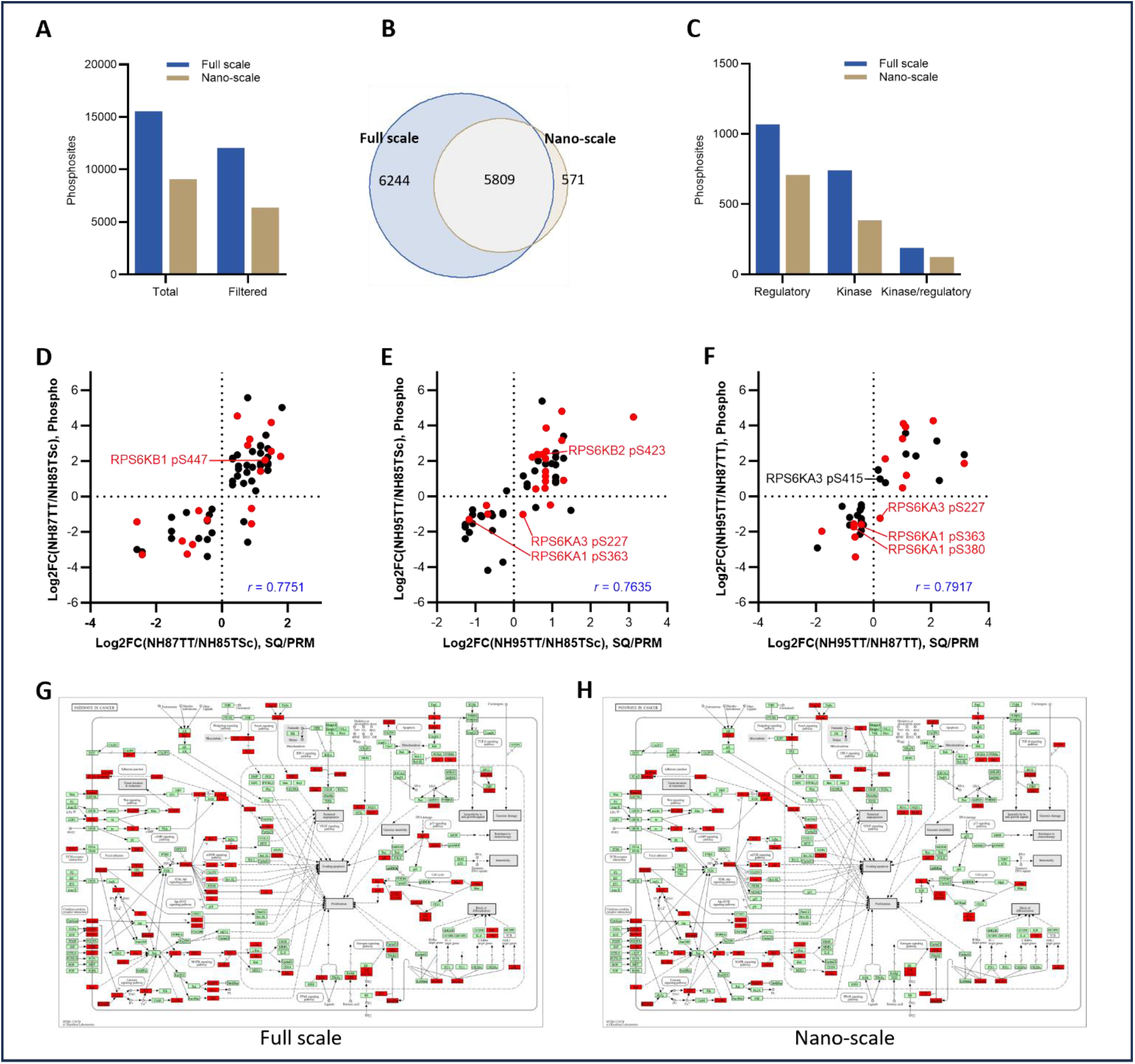
Nano-scale phosphoproteomics reveal similar findings and breadth as full-scale phosphoprotomics. Nano-scale phosphopeptide enrichment from 10 µg of tryptic peptide was performed in parallel with full scale phosphopeptide enrichment from 100 µg tryptic peptide from the same samples. **A-C)** Comparison of full-scale phosphoproteomics from B-G and nano-scale phosphopeptide enrichment using 10 µg starting tryptic peptide. Filtered phosphosites refer to sites that were quantified in all three biological replicates of at least one PDO line. **D-F)** Relative abundance of total kinase protein was plotted against the relative abundance of phosphorylation events on those kinases for each pairwise comparison between the three PDO lines. Only statistically significantly different phosphosites and kinase proteins were plotted. Red circles indicate phosphosites with defined regulatory functions. **G-H)** Comparisons of the breadth of phosphosite coverage is indicated by red coloring of nodes in the Cancer Kyoto Encyclopedia of Genes and Genomes (KEGG) pathway map05200: Pathways in Cancer.

